# Arthritis-Associated Inflammation Remodels Colonic O-Glycosylation

**DOI:** 10.64898/2026.05.20.726588

**Authors:** Piaopiao Pan, Yanling Lan, Aristotelis Antonopoulos, Stuart M. Haslam, Anne Dell, Lyin Cheng, Sai Sreeja Samavedam, Margaret M. Harnett, Simon Milling, Miguel A. Pineda

**Affiliations:** Centre for the Cellular Microenvironment, University of Glasgow, 11 Chapel Lane, G11 6EW, Glasgow, UK; Department of Life Sciences, Faculty of Natural Sciences, Imperial College London.; School of Infection and Immunity, University of Glasgow, 120 University Place, G12 8TA, Glasgow, UK

**Author notes:** Department of Wine, Vine & Beverage Sciences, School of Food Sciences, University of West Attica, Athens, Greece (Aristotelis Antonopoulos).

**Keywords:** Gut immunology, arthritis, mucins, O-glycosylation, intestinal fibroblast

## Abstract

The gut-joint axis describes how impaired intestinal epithelial function and increased gut permeability allow luminal factors to enter circulation. This can drive inflammation in Rheumatoid Arthritis, a chronic condition affecting the joint with systemic features. What mechanisms contribute to disease persistence are, as yet, incompletely understood. In health, extensively O⍰glycosylated intestinal mucins are central to epithelial protection and immune homeostasis; however, whether mucin glycosylation is altered during arthritis has not been addressed.

Here, we investigated whether arthritis⍰associated inflammation alters mucin O⍰glycosylation, potentially compromising intestinal barrier function. Using a collagen⍰induced arthritis mouse model, we combined epithelial transcriptomics, mass spectrometry-based glycomics, and imaging approaches to profile intestinal glycosylation. We identified distinct glycan remodeling in the colon, characterized by reduced fucosylation, while the ileum remained largely unaffected. In vitro studies using 3D human epithelial cultures further demonstrated that inflammatory cues, particularly from TNF⍰activated stromal cells, are sufficient to reduce epithelial fucosylation.

Together, these findings identify a stromal-inflammatory mechanism that disrupts mucin glycosylation during arthritis. Loss of colonic fucosylation emerges as a novel element of inflammatory arthritis, providing an additional mechanistic link between intestinal inflammation and fibroblast-dependent modulation of the tissue microenvironment.

## Introduction

The intestinal surface maintains stability between immunity, tolerance, and the compartmentalization of the microbiota, whilst preserving tissue functionality. One critical element of this system is the mucus layer that covers the epithelial surfaces within the luminal space. Intestinal epithelial cells, including specialised mucus⍰secreting goblet cells, are the main site of mucin⍰type O⍰glycan biosynthesis in the gut. Goblet cells synthesize and glycosylate mucins within the epithelial compartment prior to their release into the lumen. In the small intestine, mucins form a thin and dynamic layer that facilitates nutrient absorption while providing protection. Most mucins here are membrane-bound, MUC3, MUC4, MUC12, MUC13 and MUC17, forming part of the epithelial glycocalyx ^1^. In contrast, the colon is rich in secreted mucins dominated by MUC2 that assemble into a dense, two-layered mucus barrier ^2^. This structure separates the epithelium from the highly populated colonic microbiota, protecting and maintaining immune homeostasis.

Mucins are brush-like structures, where the protein backbone holds the bristles that are O-glycans. O-glycans make up most of the mucin mass and are crucial for their function. They are branched carbohydrate chains covalently linked to serine or threonine residues. Synthesis begins with the addition of N-acetylgalactosamine (GalNAc) to the protein, forming the Tn antigen, which is then elongated into various core structures (mainly cores 1–4) by the sequential addition of galactose and N-acetylglucosamine ^3, 4^. Subsequent addition of fucose and sialic acids generates further functional and structural diversity. Fucose and sialic acid content follows distinct gradients through the gastrointestinal gut, with mucin sialylation decreasing from proximal to distal gut, whereas fucosylation reaches the highest expression in the colon ^5–7^. These glycan patterns correlate with the capacity of mucins to shape distinct microenvironment functionality and physiological needs. Higher fucosylation in the colon reflects an adaptive strategy to support symbiotic microbial communities while maintaining epithelial protection and immune homeostasis via IL-22 ^8, 9^. Furthermore, many commensal bacteria express specific fucosidases to use host fucose as an energy source, establishing sustainable networks to promote the growth of beneficial microbes ^10^.

Compromise of the intestinal epithelial integrity is an early event during intestinal infection and most forms of Inflammatory Bowel Disease (IBD) ^11^. Changes in mucin composition are also linked to impaired gut function and inflammation ^12^, with increased sialylated Tn antigen and reduced fucosylated glycans reported ^13–15^. Further supporting the role of O-glycosylation in gut homeostasis, loss of core 1 O-glycans and loss of fucosylation induce colitis in mice ^16,17^, and gut dysbiosis in the microbiome ^18, 19^. Overall, these findings have led to a significant increase in research of intestinal inflammatory pathways, aiming to identify targets to manipulate gut function in local, but also systemic inflammation. When the gut barrier function weakens, bacterial and harmful substances can leak into the bloodstream and spark widespread inflammation, which contributes to systemic disease. For instance, in Rheumatoid Arthritis (RA), joint pathology is associated with gut inflammation ^20 21, 22^, with shifts in gut bacteria appearing even before joint pathology begins ^23^. Such changes have been also associated with disease phenotypes ^24^. Collectively, these findings shaped the concept of the ‘gut-joint axis’, a mechanism postulated to link loss of gut permeability, with microbial and immune dysregulation, that may contribute to the inflammatory disease in the joint ^25^.

Experimental and clinical studies have been conducted to support the gut-joint axis, aiming to identify key molecular mechanisms and therapeutic targets. However, despite the prevalent role of O-glycans in the gut, a study defining the intestinal O-glycome during arthritis has not been conducted. Herein, we hypothesised that the inflammatory process underlying arthritis remodels the O-glycome in the gut, which could subsequently disrupt local homeostasis. We have focused on intestinal epithelial cells, as the primary site of mucin⍰type O⍰glycan biosynthesis in the gut, expressing the glycosylation enzymes required for O⍰glycan assembly of all secreted and membrane⍰associated mucins. Experimentally, we utilised healthy mice and mice undergoing collagen-induced arthritis (CIA). Gut tissue was dissected at peak of disease to conduct i) transcriptomic profile of O-glycan biosynthetic pathways in epithelial cells, and ii) mass-spectrometry based glycomics of isolated O-glycans. Our results show that arthritic mice reduce fucosylation in the colon, increasing relative sialylation, whereas the O-glycome in the ileum remains unaltered. Finally, we conducted *in vitro* experiments in 3D gut models showing that loss of epithelial fucosylation can be reproduced by TNF-mediated stimulation of gut fibroblasts. Our findings reveal a previously unappreciated connection between inflammatory mediators and colonic O⍰glycosylation, supporting a role for altered mucosal glycosylation in the gut-systemic inflammatory axis.

## Results

### Arthritic mice show altered gut morphology and tissue damage

We chose collagen-induced arthritis (CIA) as our experimental model for inflammatory arthritis. In this model, joint inflammation is induced by injection of type II collagen, which induces strong IL-17/IL-1β responses. Around day 30-35, joints exhibit evident signs of disease and pathophysiological responses characteristic of human disease, including the infiltration of immune cells, the production of autoantibodies, and the activation of synovial fibroblasts, leading to the formation of pannus (Supplementary Figure 1) ^26^. CIA mice have also exhibited gut inflammation, intestinal damage, and pathogenic dysbiosis ^21, 22^, mirroring the gut-joint pathogenic axis proposed in human disease. Our results here further corroborate this, arthritic mice showed irregular villi in the ileum (Figure 1A) and loss of consistent mucin expression and normal tissue architecture in the colon (Figure 1B). Whereas colon from naïve tissue displays well-organised crypt architecture with uniform goblet cell distribution and consistent mucin staining, arthritic colon exhibited mild crypt disorganisation and heterogeneous Periodic acid-Schiff (PAS) staining, indicative of altered mucin production. No major epithelial erosion or severe inflammatory infiltration was observed at the histological level (Figure 1B). These results indicate that colonic alterations in arthritic mice may be associated with mucosal remodelling, reduction in goblet cell distribution and disruption of mucin homeostasis. Such observations provided additional support to our working hypothesis, which placed mucin glycosylation as a key factor in the pathophysiology of arthritis.

**Figure 1.**
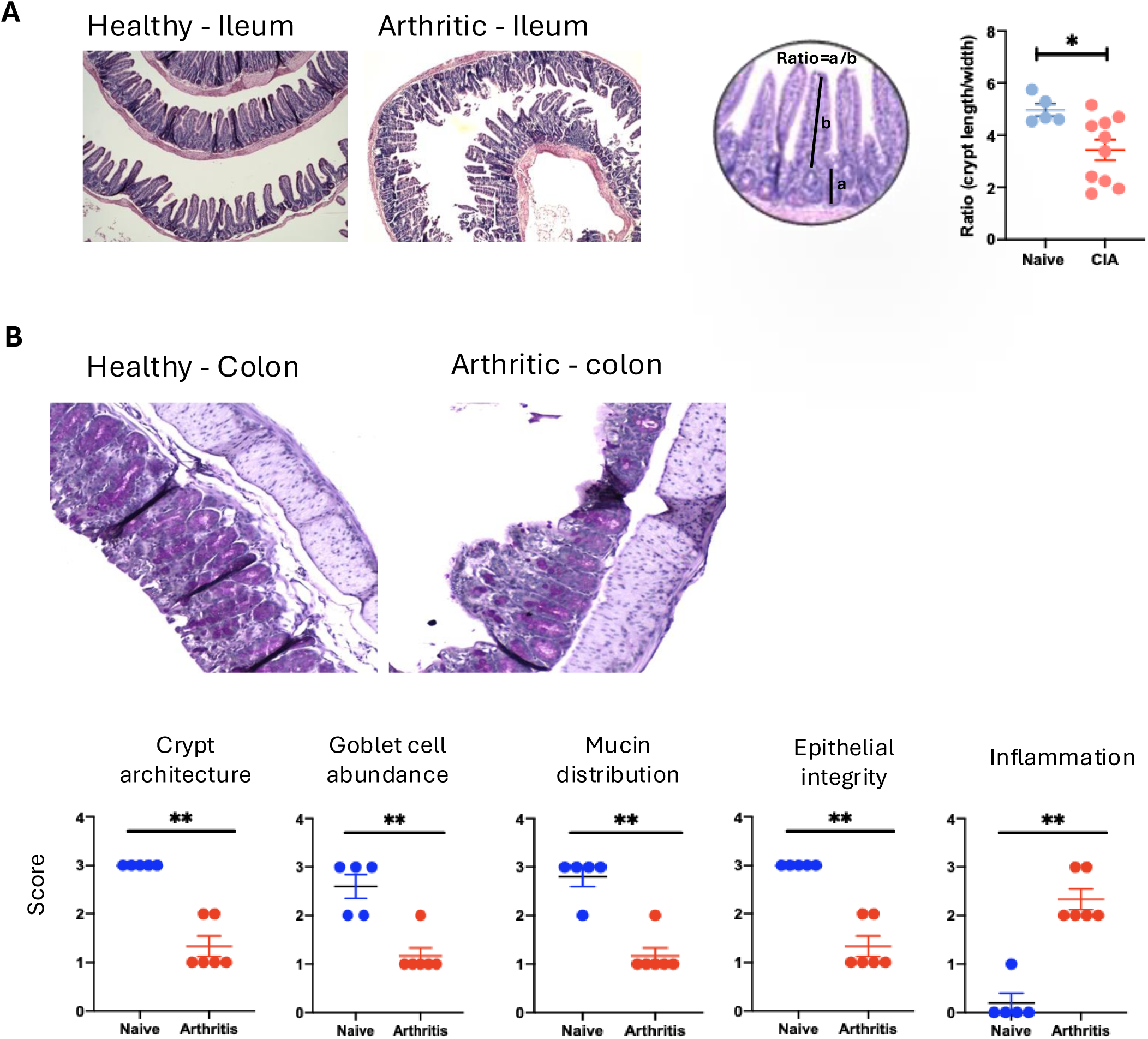
Intestinal histopathology shows ileal and colonic mucosal damage in arthritic mice. (A) Representative hematoxylin and eosin (H&E) stained sections of ileum from healthy (naïve) and collagen⍰induced arthritis (CIA) mice. Arthritic mice show marked disruption of ileal architecture, characterized by villus blunting and shortening. Villus morphology was quantified as the ratio of villus length (a) to villus width (b), as illustrated in the micrograph. Quantitative analysis in the graph shows the villus length⍰to⍰width ratio in arthritic mice compared with naïve controls. (B) Representative periodic acid-Schiff (PAS) stained sections of colon from naïve and arthritic mice, highlighting goblet cells and mucin content. Colonic tissue from arthritic mice exhibits architectural distortion of crypts, reduced goblet cell abundance, altered mucin distribution, compromised epithelial integrity, and increased inflammatory infiltrates. Bottom panels show semi⍰quantitative histopathological scores for crypt architecture, goblet cell abundance, mucin distribution, epithelial integrity, and inflammation. Scoring system goes from 0 (low) to 3 (high) in each case. Each dot represents one individual mouse, with mean ± SEM. Statistical significance was determined using unpaired t test, with ** = p < 0.01.

### Ileum and colon present distinct glycosylation signatures

Glycans in gut mucins play distinct biological roles in different intestinal regions. In the ileum, they contribute to antigen sampling and the initiation of adaptive immune responses, whereas in the colon they are involved in maintaining tolerance and diversity of commensal microbiota. Accordingly, the ileum and colon were analysed separately in our experiments. To obtain an overall view of tissue glycosylation, we first selected plant lectins that recognise glycan motifs relevant to epithelial mucins, covering fucose and sialic acid-containing glycans, that in mice are mostly associated to core-1 and core-2 structures (Figure 2A): i) UEA recognises the motif fucose α1-2Galactose; ii) PNA recognises the T-antigen, or core-1 structure Galβ1-3GalNAc, only when it is not sialylated, and iii) MAL-II and SNA recognise sialic acid in α2-3 and α2-6 glycosidic linkages respectively. As previously reported, the colon of healthy animals exhibited higher levels of fucosylation and sialylation than the ileum, as indicated by increased UEA, MAL-II, and SNA binding (Figure 2B). In contrast, non-sialylated T-antigen levels were reduced in the colon, as shown by decreased PNA binding (Figure 2B). Consistently, fucose and sialic acid binding lectins showed a strong colocalization with MUC2 in the colon, the major mucin produced by goblet cells (Supplementary Figure 2). MUC2 fucosylation was found in the luminal and intercrypt mucus. Sialylated MUC2 showed distinct spatial patterns, with α2,6-sialylated mucins expressed more broadly across the tissue that α2,3-linkages. In contrast, PNA staining for exposed Galβ1-3GalNAc core structures showed limited colocalization with MUC2, consistent with masking of core O-glycan structures in mature mucus by terminal sialylation that blocks PNA binding. These results demonstrate tissue- and site-specific glycosylation of gut mucins.

**Figure 2.**
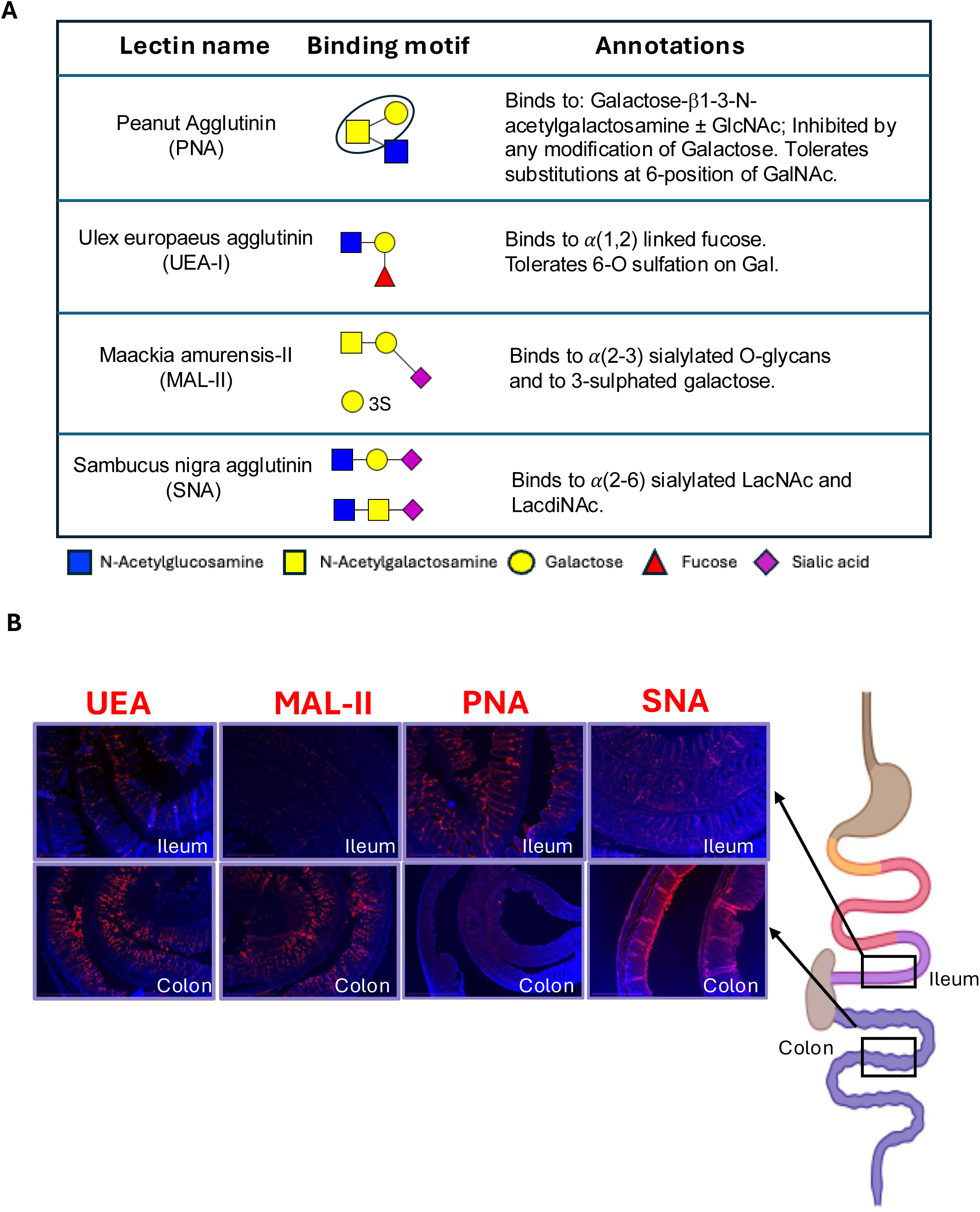
Lectin⍰based glycan profiling reveals regional distribution of carbohydrate motifs in the gut of naïve mice. (A) Summary of lectins used for histological labelling, including their canonical glycan⍰binding specificities and structural motifs. Peanut agglutinin (PNA) recognizes Galβ1⍰3GalNAc; Ulex europaeus agglutinin I (UEA⍰I) binds α(1,2)⍰linked fucose; Maackia amurensis lectin II (MAL⍰II) recognizes α(2,3)⍰sialylated O⍰glycans; and Sambucus nigra agglutinin (SNA) binds α(2,6)⍰sialylated LacNAc structures. Symbol key indicates monosaccharide composition. (B) Representative fluorescence images of lectin staining in ileal and colonic sections from naïve mice. Differential labelling by UEA⍰I, MAL⍰II, PNA, and SNA reveals region⍰specific distribution of fucosylated and sialylated glycans along the intestinal epithelium and within the mucus layer. Schematic indicates anatomical location of sampled ileal and colonic regions. Nuclei are counterstained with DAPI (blue). Scale bars, 500 µm.

### Inflammatory arthritis down-regulates fucosylation biosynthetic pathways in colonic epithelial cells

Once the distinct fucosylation and sialylation patterns were confirmed in ileum and colon of healthy tissue, we next aimed to identify changes in epithelial glycan expression during arthritic conditions. First, we isolated intestinal epithelial cells to perform RNA sequencing, to define transcriptional programs underlying O-glycosylation in naïve and arthritic mice. Following tissue digestion, cells were sorted from the epithelial compartment as CD31⁻CD45⁻EpCAM⁺ (Figure 3A). This population predominantly includes enterocytes, goblet cells, and Paneth cells in the ileum, and mainly enterocytes and goblet cells in the colon. The percentage of isolated epithelial cells in the total tissue was approximately 10% in the ileum and 5% in the colon, with no changes observed in arthritic mice (Figure 3B). An initial non-targeted analysis reveals activation of pathogenic and inflammatory pathways in the colon, but not ileum, of arthritic mice. PCA analysis of all transcripts showed clustering (Figure 3C) and differential gene expression in the colon (Figure 3D), with 70 differentially expressed genes in arthritic colon [27 up-regulated and 43 down-regulated; >2-fold, padj□<□0.01] (Figure 3E). The up-regulated genes primarily represent an inflammatory signature characteristic of immune cells rather than epithelial cells. However, the total transcript counts for these genes are very low (Figure 3F), suggesting that this signal may reflect ambient contamination from RNA released by nearby immune cells during tissue processing, low-level transcription, or technical artefacts during cell sorting rather than active expression in epithelial cells. In contrast, the down-regulated genes display a strong epithelial signature, with high transcript abundance (Figure 3F). Down-regulated genes indicated that epithelial cells in arthritic colon exhibit reduced secretory capacity [SPDEF,TFF2,PLET1,UPK1B] attenuated stress-response pathways [LAMB1, P4HA2, ADAMTS17, CCN3, BCAN], and diminished extracellular matrix interactions [HSPB1, HSPH1, MT3] (Figure 3E).

**Figure 3.**
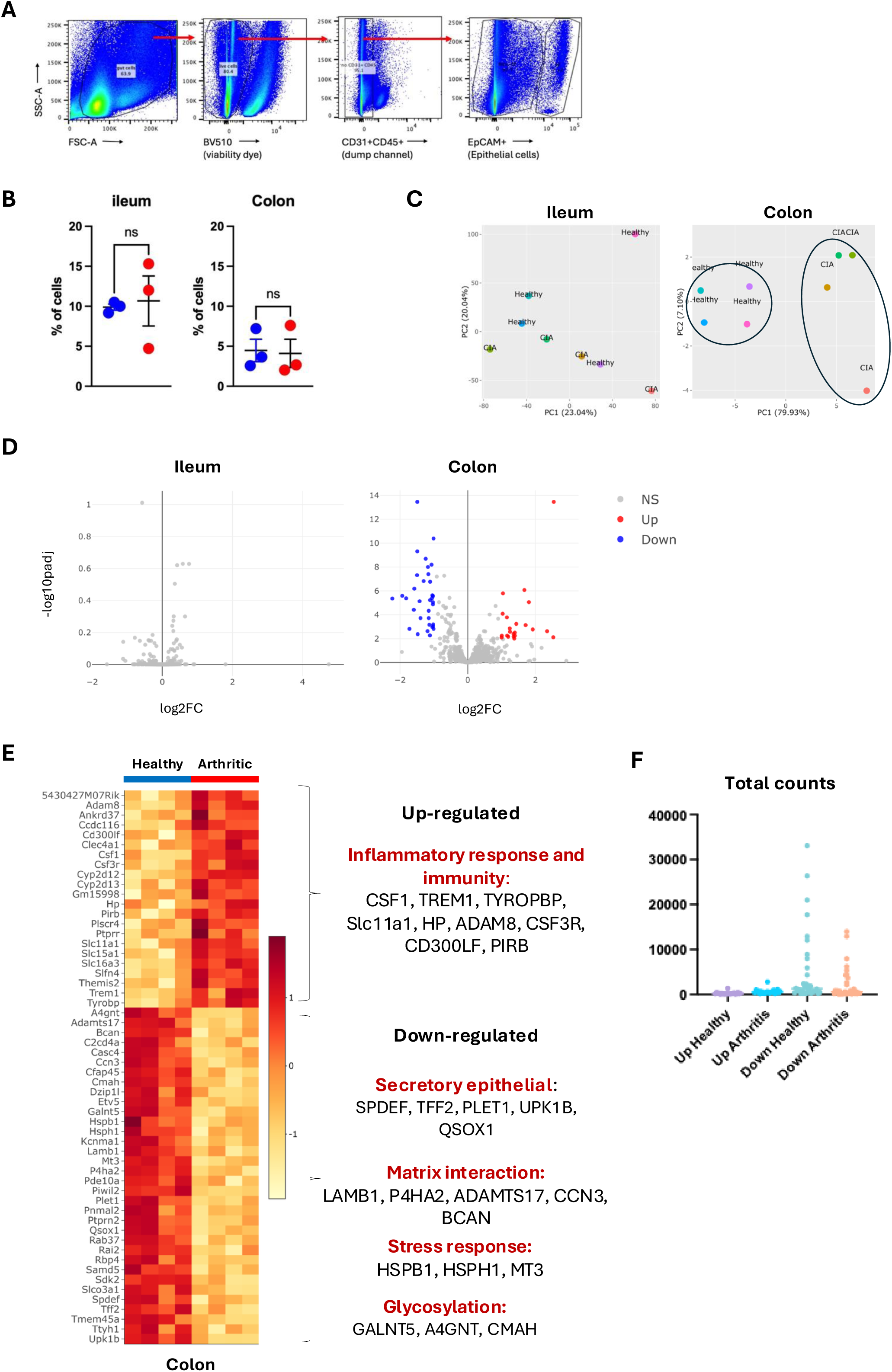
Transcriptomic profiling of intestinal epithelial cells reveals disease⍰ and region⍰specific alterations in arthritic mice. Ileum and colon tissues were collected from healthy and collagen-induced arthritis (CIA) mice and digested ex vivo to isolate epithelial cells. (A) Epithelial cells were identified by flow cytometry as live cells EpCAM+ and lacking CD31 and CD45 (markers for endothelial and immune cells). (B) Relative numbers of epithelial cells are shown for individual mice (blue circles, healthy; red circles, arthritic). Statistical significance was assessed by one-tailed t-test; ns = not significant. (C) Principal component analysis (PCA) of RNA⍰seq data from ileal and colonic epithelial cells demonstrates tissue⍰specific clustering and distinct transcriptional profiles between naïve and arthritic mice in the colon, with minimal separation in the ileum. (D) Volcano plots depicting differentially expressed genes (DEGs) analysed by DeSeq2 [adjp < 0.01, fold difference > 2] between naïve and arthritic mice in ileal and colonic epithelial cells. Red and blue dots indicate significantly up⍰regulated and down⍰regulated genes, respectively; non⍰significant genes are shown in grey. (E) Heatmap of DEGs in colonic epithelial cells highlighting up⍰regulated inflammatory and immune⍰associated pathways and down⍰regulated genes involved in epithelial secretory function, extracellular matrix interactions, stress responses, and glycosylation in arthritic mice compared with naïve controls (n=4 per group). (F) Total read counts for DEG genes across all samples in arthritic colon epithelial cells. Each dot represents one specific gene.

However, relatively few genes were differentially expressed in the global RNA⍰seq analysis. Several genes involved in glycosylation (including Galnt5, A4GNT, and CMAH) were detected in the down⍰regulated group, suggesting alterations in glycosylation-related pathways. Mucin O⍰glycosylation is governed by coordinated activity of multiple glycosyltransferases and modifying enzymes, often with modest transcriptional changes, and standard bioinformatic pipelines fail to capture coordinated transcriptional regulation across O-glycosylation pathways. Thus, we performed a targeted, pathway⍰focused analysis of mucin glycosylation genes within the RNA⍰seq dataset. This biologically informed approach increases sensitivity for detecting coordinated transcriptional shifts across key components of the O⍰glycan biosynthetic machinery that may not reach significance in transcriptome⍰wide analyses, thereby providing clearer mechanistic insight into how inflammation impacts epithelial mucin composition. A curated list of genes to study mucin O-glycosylation was created (Figure 4A) to include glycosyltransferases involved in O-glycan synthesis: i) Polypeptide N-acetylgalactosaminyltransferases (ppGalNAcTs; transfer of N-acetylgalactosamine (GalNAc) from UDP-GalNAc to serine or threonine residues), ii) C1GalT1 and GcnTs, for core 1-4 formation, iii) sialyltransferases (ST3GalTs and ST6GalNAcTs) for addition of sialic acid and iv) fucosyltrasferases, FutTs, responsible for fucose incorporation. Using only this gene list, we conducted a new PCA to evaluate whether epithelial cells in arthritic mice exhibited differences in O-glycosylation-related transcriptional programs. No significant differences were observed in the ileum (Figure 4B), whereas in the colon, the O-glycosylation biosynthetic signature did clearly separate naïve and arthritic mice (Figure 4C). Next, we examined individual glycosylation-related genes, using both Z-scores and total normalised counts, to highlight which enzymes contribute most to the O-glycosylation signature in each tissue, in healthy and inflammatory conditions (Figure 5). The ileum had low expression levels of sialyl- and fucosyltransferases (Figure 5A), in line with the lectin-based results seen before (Figure 2). No significant changes were observed in the ileum of arthritic mice, but data obtained from colon (Figure 5B) showed a different scenario. Overall, sialyl- and fucosyltransferases showed higher expression in the colon compared with the ileum in healthy tissue, and O-glycosylation-related gene expression patterns distinguished naïve and arthritic groups. Most Galnt genes, except Galnt1, were significantly down-regulated in arthritis, as were the core 1 forming enzymes C1GalT1 and Cosmc, consistent with a potential reduction in overall mucin O-glycosylation. Terminal fucosylation is mainly mediated by Fut2 and Fut4, which were both highly expressed and significantly downregulated in arthritic mice, while other FUT family members showed very low or undetectable expression. Regarding sialylation, only ST3Gal3 was significantly downregulated in arthritis, whereas other enzymes did not reach any statistical difference. This transcriptomic profile of arthritic colon epithelial cells suggests a reduction in mucin O-glycosylation capacity, that in conjunction with significant downregulation of Fut2 and Fut4, points to globally reduced fucose content. Sialylation may also be subtly altered, but there was no consistency among enzymes to predict functional impact in sialylated glycan synthesis. Collectively, these changes predict colonic mucins with lower glycan density and fewer terminal fucose substitutions.

**Figure 4.**
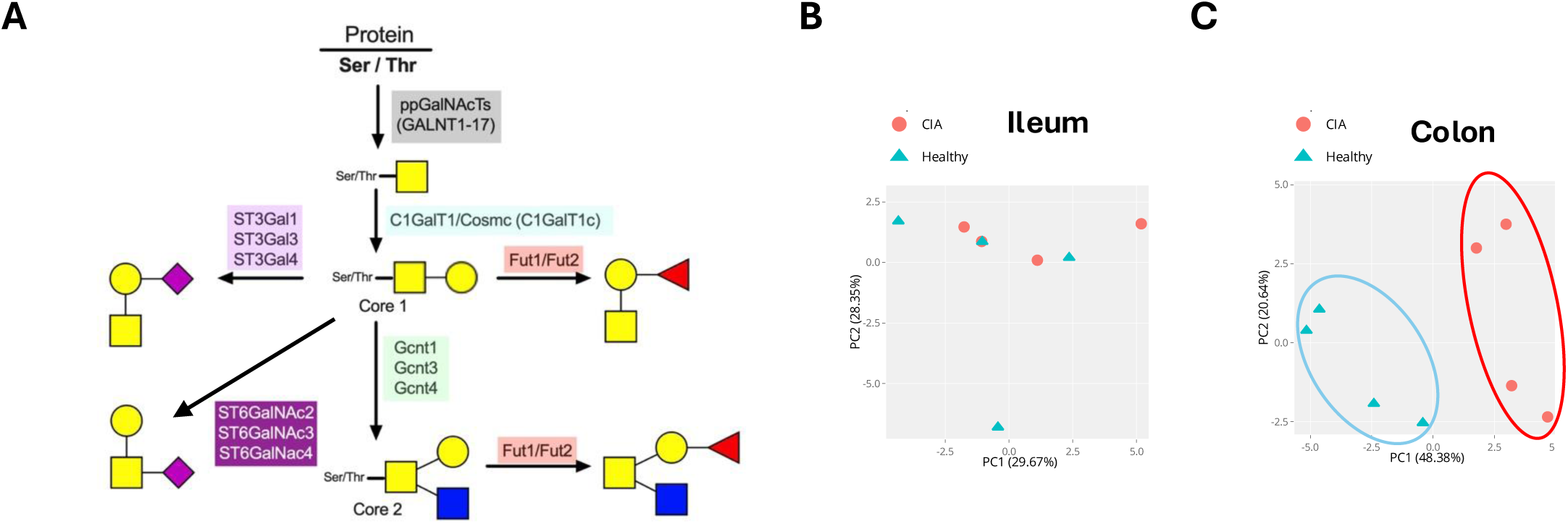
Targeted analysis of mucin O⍰glycosylation pathway gene expression in intestinal epithelial cells from naïve and arthritic mice. (A) Schematic overview of the mucin⍰type O⍰glycosylation pathway in murine mucins, illustrating initiation by polypeptide N⍰acetylgalactosaminyltransferases (ppGalNAcTs; GALNT1–17), formation of core 1 and core 2 O⍰glycan structures by C1GALT1/Cosmc and GCNT family enzymes, and subsequent terminal modification by sialyltransferases (ST3Gal, ST6GalNAc) and fucosyltransferases (Fut1/Fut2). (B, C) Principal component analysis (PCA) of RNA⍰seq data restricted to genes involved in mucin O⍰glycosylation in epithelial cells isolated from the ileum (B) and colon (C) of naïve and collagen⍰induced arthritis (CIA) mice. Each dot represents an individual mouse (blue, healthy; red, arthritic).

**Figure 5.**
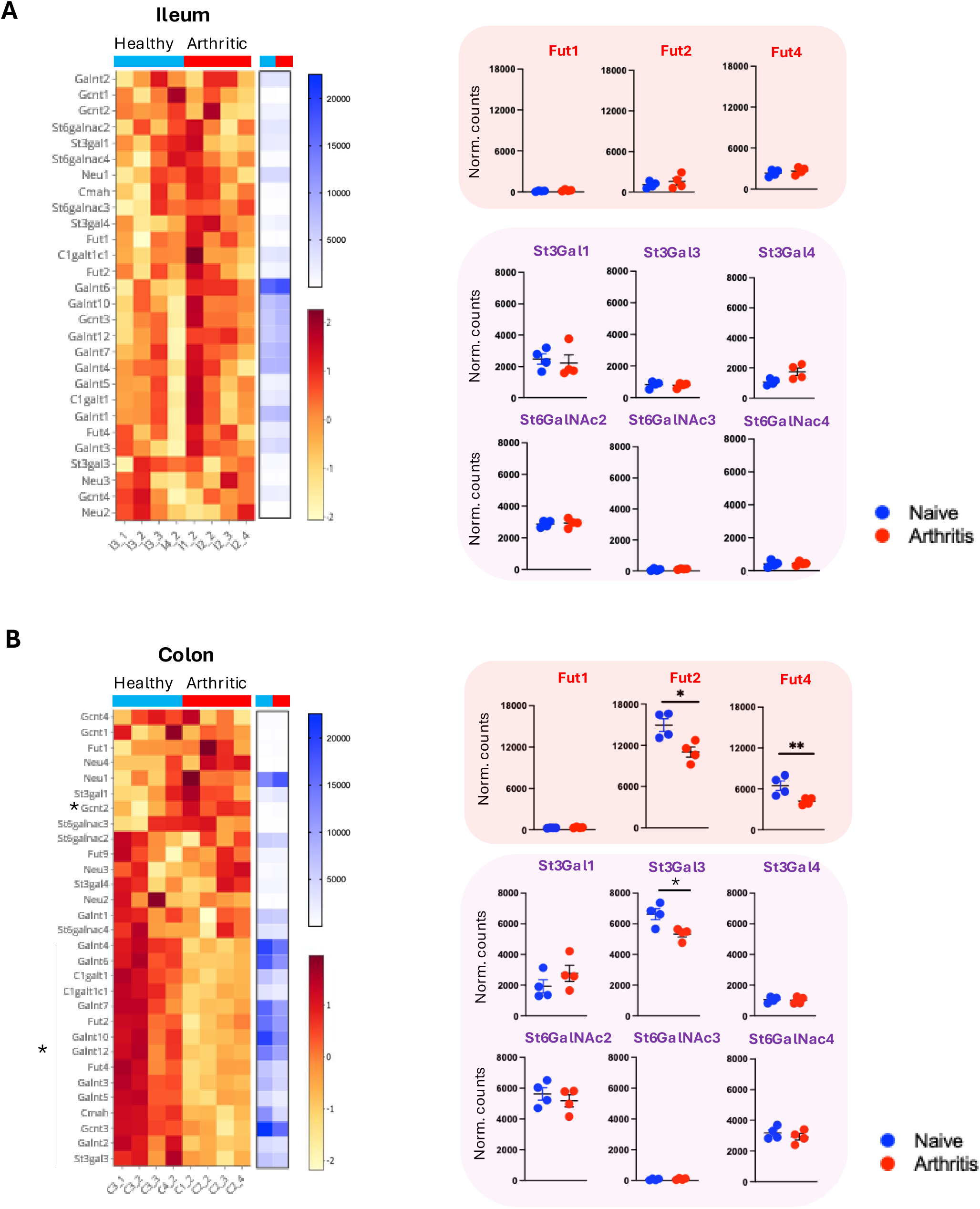
Gene⍰level alterations in mucin O⍰glycosylation pathways in intestinal epithelial cells from naïve and arthritic mice. (A) Heatmap showing expression of genes involved in mucin⍰type O⍰glycosylation in ileal epithelial cells isolated from naïve and collagen⍰induced arthritis (CIA) mice. Gene expression is displayed as z⍰scores calculated across samples for each gene and visualized using a red-yellow colour scale. Adjacent blue-white heatmap indicates normalized RNA⍰seq counts for the same genes. Right panels show normalized counts for detected fucosyltransferases (Fut1, Fut2, Fut4) and detected sialyltransferases (St3gal1, St3gal3, St3gal4, St6galnac2, St6galnac3, St6galnac4) plotted for individual mice. (B) Corresponding analysis of colonic epithelial cells. Heatmaps depict z⍰score–scaled expression (red-yellow) and normalized counts (blue-white) for mucin O⍰glycosylation genes, with graphical representation of normalized counts for the same selected enzymes shown on the right. Colonic epithelial cells from arthritic mice exhibit pronounced disease⍰associated changes in expression of fucosylation⍰ and sialylation⍰related genes compared with naïve controls. Data points represent individual mice; bars indicate mean ± SEM. Statistical significance, unpaired t-test, is indicated where applicable (*p < 0.05).

### Arthritis increases the fucose/sialic acid ratio in the colonic O-glycome

While transcriptomic analysis of glycosyltransferases provides insight into the regulatory programs governing glycosylation, gene expression does not directly reflect the structure or abundance of glycans, which are shaped by additional factors such as enzyme activity, substrate availability, and commensal microbiota. To directly assess the functional outcome of these transcriptional changes, we performed mass spectrometry-based studies to characterize the glycan profiles of ileal and colonic tissue, focusing on O-glycans as a proxy for mucin glycosylation under naïve and arthritic conditions. Based on the observed down-regulation of key glycosyltransferases, including those involved in glycan initiation and terminal modification, arthritic tissues could display reduced O-glycan complexity and reduced terminal fucosylation. O-glycans were released by reductive elimination and permethylated, prior to mass-spectrometry analysis for further characterisation and quantification. Specific mass (*m/z*) peaks for well-described motifs in murine mucin O- glycans were evaluated in ileum and colon (Figure 6A). The colon showed a more diverse glycome, with high expression of sialylated and fucosylated core 1 and core 2 structures. Peak intensity for specific glycans were evaluated in naïve and arthritic mice, including non-modified core structures [core 1, core 2] (Figure 6B), fucosylated-core 1 which showed a statistically significant decrease (Figure 6C), and sialylated-core 1 which showed a statistically significant increase (Figure 6D). Overall, arthritic mice exhibit a remodelling of colonic O-glycans, characterised by the physiological fucosylation of core structures being redirected towards an abnormal sialylated profile. This phenomenon is specific to the colon, as no alterations are observed in the O-glycan repertoire of the ileum.

**Figure 6.**
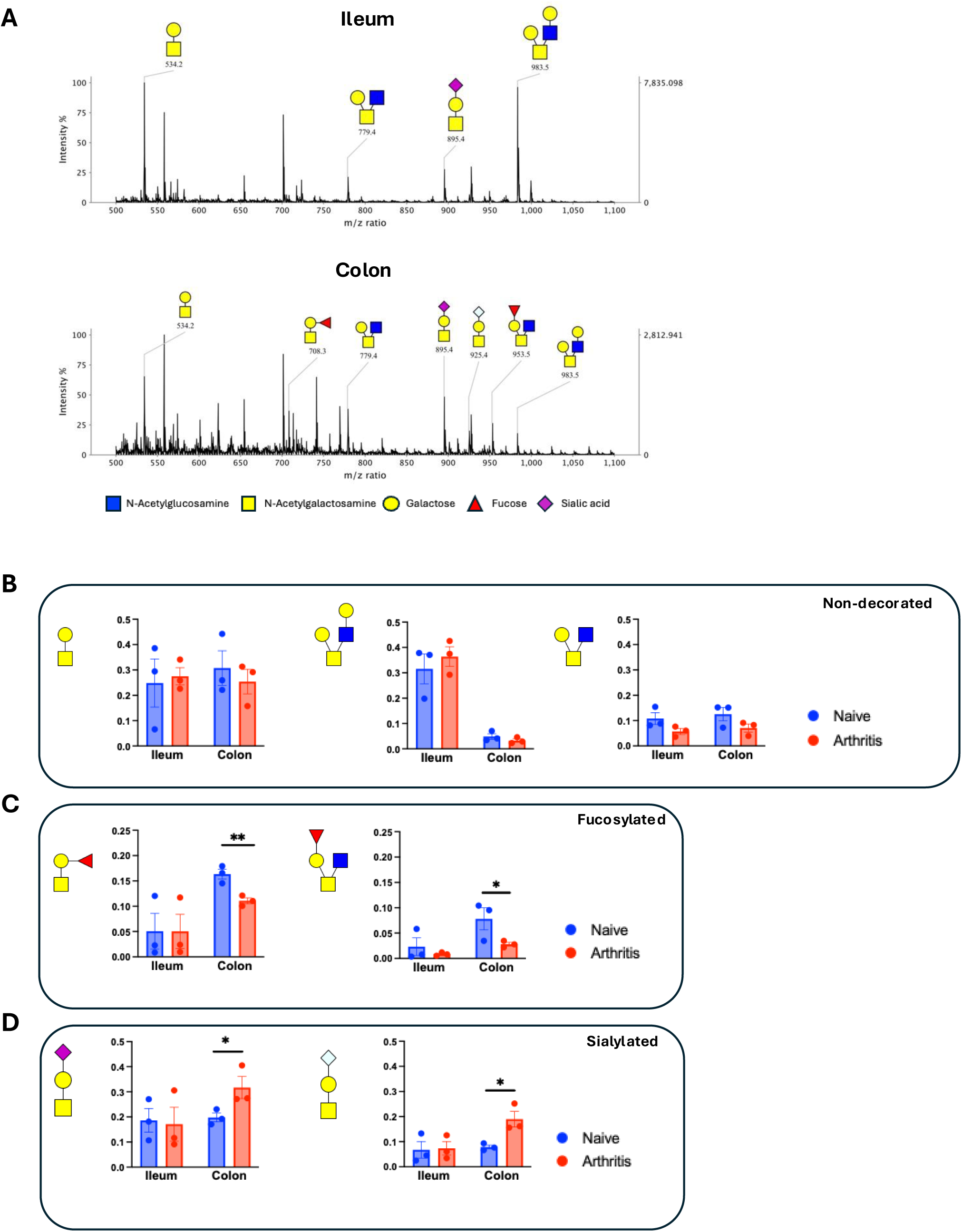
Mass spectrometry–based glycomic profiling reveals region⍰ and disease⍰specific alterations in colonic O⍰glycan structures. (A) Representative MALDI⍰TOF mass spectra of O⍰linked glycans released from ileal and colonic tissue of naïve mice. Annotated peaks indicate major glycan species, with monosaccharide compositions schematically represented. Symbols denote N⍰acetylglucosamine (blue square), N⍰acetylgalactosamine (yellow square), galactose (yellow circle), fucose (red triangle), and sialic acids (diamonds, purple: 5NeuAc, blue: 5NeuGc). (B) Relative abundance of selected non⍰decorated core-1 and core-2 O⍰glycan structures in ileum and colon from naïve (blue bars) and collagen⍰induced arthritis (CIA) mice (red bars). Data are shown as proportion of total detected O⍰glycans. (C) Quantification of fucosylated O⍰glycan species in ileal and colonic samples. (D) Quantification of sialylated O⍰glycan species in ileum and colon. For B-D, individual data points represent individual mice; bars indicate mean ± SEM. Statistical significance is indicated where applicable (*p < 0.05, **p < 0.01), by one-tailed unpaired t test.

### Paracrine signalling from TNF-activated fibroblasts disrupts epithelial barrier function and fucosylation

Our *in vivo* findings demonstrate significant O-glycan remodelling in the colon of arthritic mice associated with tissue damage. However, the underlying mechanisms responsible for this remodelling is not known. To investigate pathophysiological mechanisms, we worked with a 3D model of human gut epithelium. To build the model, a thin layer of collagen was first added on top of 3D porous scaffolds to facilitate the formation of a epithelial layer. Caco-2 cells were used to represent colon epithelial cells, and HT29 Goblet cells were added to the culture (10:1 Caco-2 : HT29 ratio) to recreate a functional mucus layer. Histology of scaffold sections and electron microscopy confirmed the formation of a homogenous epithelium (Figure 7A). For a functional readout for barrier integrity, we measured the flux of horseradish peroxide (HRP) from top to bottom chambers. The 3D model prevented HRP leakage under non-inflammatory conditions, but exposure to TNF disrupted this and allowed HRP translocation through the cell monolayer and scaffold, mimicking inflammation-derived gut damage and validating the model (Figure 7B). The presence of a fucosylated mucus was confirmed by UEA binding (Figure 7C). However, despite TNF induced loss of barrier permeability, TNF did not affect fucose content, suggesting that glycan remodelling is not driven directly by TNF-stimulation of epithelial cells (Figure 7C). Because TNF is a major inflammatory driver in arthritis, we hypothesised that some intermediary was required to link TNF effects and loss of O-glycan fucosylation. We turned our attention to gut fibroblasts, since their activation has been recognised as a key driver of epithelial barrier breakdown in intestinal inflammation ^27^. Fibroblasts actively shape inflammatory responses through cytokine production, ECM remodelling, and crosstalk with epithelial cells. We used the CCD-18Co human colon fibroblast cell line as a model. CCD-18Co cells were activated *in vitro* with TNF, and the conditioned medium was collected for subsequent stimulation of Caco2–HT29 epithelial layers. We first confirmed fibroblast activation, as TNF stimulation induced high levels of IL-6 and MMP3 production in culture supernatants (Figure 7D). This fibroblast-conditioned medium was then applied to 3D epithelial layers to assess lectin binding (Figure 7E). Fibroblast TNF-conditioned medium disrupted epithelial glycosylation, reducing overall fucosylation while concentrating remaining UEA binding into patchy, likely goblet cell-associated regions. Image quantification confirmed the reduced fucose expression in response to TNF-conditioned fibroblast medium (Figure 7F). Effect seemed to be specific to fucosylation pathways, since PNA binding was not affected, suggesting a lack of changes in T-antigen or sialic acid expression in O-glycans. Together, these findings identify a TNF-fibroblast-fucosylation axis that modulates the epithelial glycome in the colon upon systemic inflammation.

**Figure 7.**
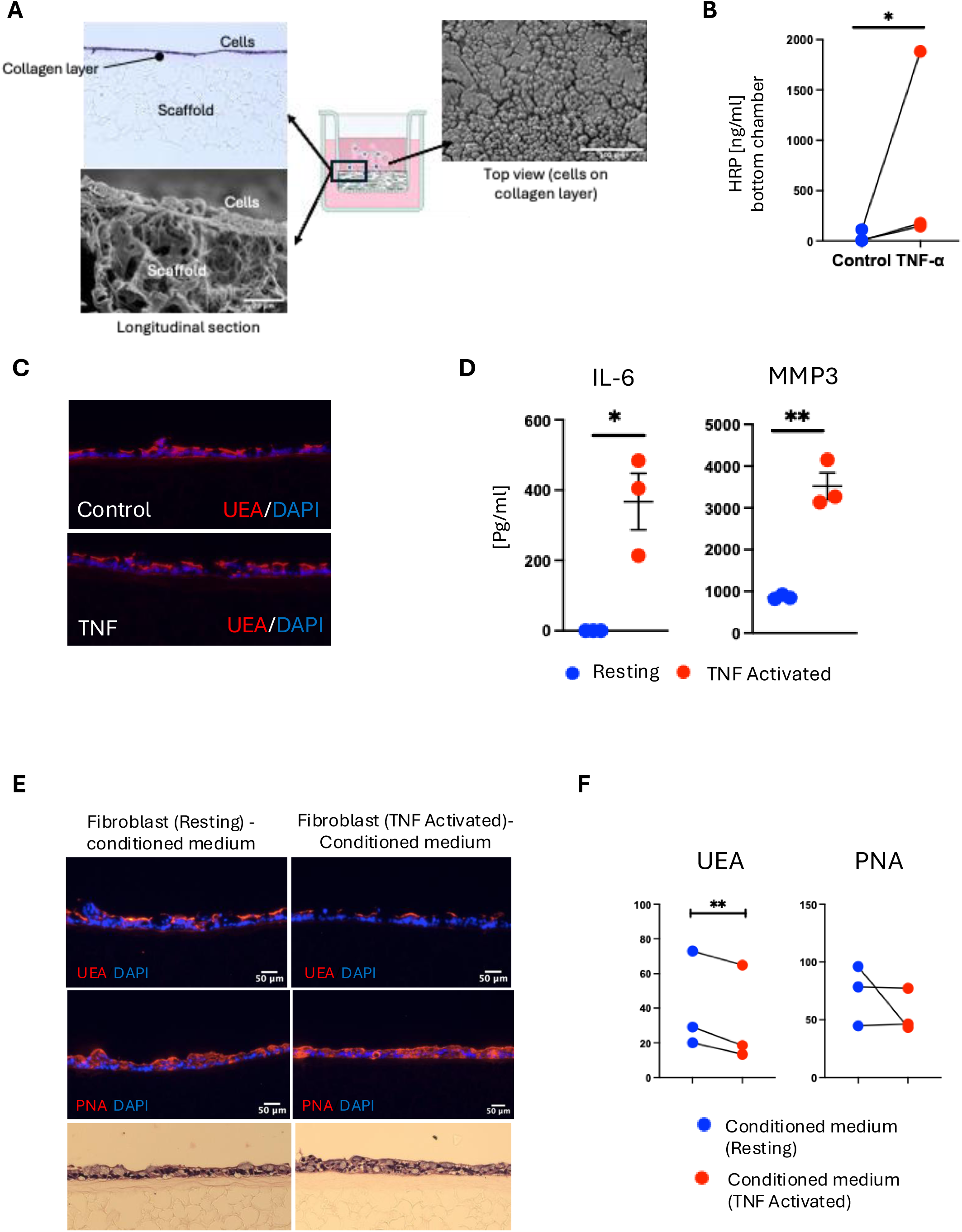
Fibroblast⍰derived inflammatory signals modulate epithelial glycan presentation. (A) 3D colon epithelial models were generated using Caco-2 (enterocyte) and HT29 (goblet) cell lines cultured on collagen-coated 3D scaffolds. H&E staining and electron microscopy confirmed epithelial organisation and formation of a continuous monolayer. (B) Barrier integrity of 3D epithelial cultures was assessed following TNF stimulation (10ng/ml, 24 hours) by measuring HRP translocation from the apical to the basal compartment. (C) UEA lectin staining of 3D cultures revealed modulation of O-glycan terminal structures in response to TNF. (D) Gut fibroblasts (CCD-18Co) were stimulated with TNF, and IL-6 and MMP3 production was quantified in supernatants by ELISA. Each dot represents one independent experiment performed in technical triplicates. *p < 0.05, **p < 0.01 by one-tailed unpaired t-test. (E) Representative immunofluorescence images of 3D epithelial layer cultured with conditioned medium from resting fibroblasts or TNF⍰activated fibroblasts. Cells were stained with UEA⍰I lectin to detect α1,2⍰fucosylated glycans (red), or peanut agglutinin (PNA; Galβ1⍰3GalNAc, red) and counterstained with DAPI (blue). H&E staining of the corresponding monolayers are shown. (F) Quantification of UEA⍰I and PNA staining intensity in epithelial monolayers cultured with conditioned medium from resting or TNF⍰activated fibroblasts. Data are shown as paired measurements for matched experiments analysed in triplicate. Statistical significance was determined using paired statistical tests; p□<□0.01.

## Methods

### Mice and induction of Collagen-Induced Arthritis (CIA)

Male DBA/1 mice (8–10 weeks; Envigo, UK) were housed under specific pathogen-free conditions at the University of Glasgow with *ad libitum* access to food and water. Animals were randomly assigned to experimental groups, and all assessments were performed by investigators blinded to group allocation. All procedures adhered to ARRIVE guidelines and were approved by the University of Glasgow AWERB and carried out under UK Home Office licences (PPL P8C60C865, PPL PE1AF6E8B, PPL PP9710134; PIL I62988261, I675F0C46). CIA was induced by intradermal injection at the tail base of 100 μg chicken type II collagen (CII; MD Biosciences) emulsified 1:1 in complete Freund’s adjuvant (day 0). A booster of 200 μg CII in PBS was administered intraperitoneally on day 21. Mice were monitored every 2 days for body weight, paw thickness, and clinical scores (0 to 4 per paw, being 4 the highest inflammatory stage). Humane endpoints included: total clinical score >10, >20% body-weight loss, paw thickness >4.5 mm, or inflammation in ≥3 paws. Animals reaching endpoints were humanely euthanised.

### Histology

Gut tissue was fixed in 10% neutral-buffered formalin, paraffin-embedded, and sectioned at 7 μm using a Leica RM2125 microtome. Sections were incubated at 60°C, dewaxed, rehydrated through graded alcohols, and stained with Harris haematoxylin and eosin using standard protocols. Slides were dehydrated, cleared in xylene, and mounted in DPX. Brightfield images were acquired using an EVOS microscope. Villus length and crypt depth were measured in ImageJ. Colonic tissues were fixed in 10% neutral-buffered formalin, paraffin-embedded, sectioned at 7 µm, and stained with periodic acid–Schiff (PAS) using standard protocols. A semi-quantitative scoring system was applied to assess mucosal structure and inflammation, adapted from established murine colitis scoring methods ^28^. Five parameters were evaluated: crypt architecture, goblet cell abundance, mucin distribution across tissue, epithelial integrity, and inflammatory infiltrate. Each parameter was scored on a scale of 0–3.

### Gut tissue digestion

Gut digestion was performed following removal of mesenteric fat and Peyer’s patches, intestines were opened longitudinally, rinsed in PBS, and cut into 1-cm pieces. Tissues were washed in warm HBSS without Ca²⁺/Mg²⁺ containing 2 mM EDTA (37°C, 220 rpm, 15 min each). Ileal segments were digested in R10 medium containing 0.5 mg/mL collagenase IV; colonic segments were digested in R10 containing 0.5 mg/mL collagenase IV and 24 μg/mL DNase I (37°C, 220 rpm; 15 min for ileum and 20 min for colon). Digests were diluted in ice-cold RPMI, passed through 100-μm then 70-μm strainers, centrifuged (400 g, 10 min), washed in ice-cold R10, and prepared for flow cytometry.

### Flow cytometry and cell sorting

Single-cell suspensions from digested gut tissue were resuspended in PBS without serum (1-10 × 10□ cells/mL) and stained with BV510 viability dye (30 min, 4°C). Cells were washed, incubated with anti-mouse CD16/32 to block Fc receptors (15 min, RT), and stained with the antibody panel for detection of CD31-CD45-EpCAM+ epithelial cells: (CD31-AF700 [Biolegend], CD45-AF700 [eBioscience], EpCAM-FITC [Biolegend]). Following staining, cells were washed in FACS buffer (PBS, 1% FBS, 0.4% EDTA) and fixed using fixation buffer (Invitrogen; 15 min, RT). For cell sorting, samples were acquired on a FACSAria III (BD) using a 100-μm nozzle. Fractions were collected into 1.5-mL tubes containing 500 μL RLT buffer (Qiagen) and kept on ice for RNA extraction following manufacturer’s instructions.

### RNA sequencing and bioinformatic analysis

Total RNA from gut tissue and intestinal epithelial cells was extracted and quality-checked using Nanodrop (260/280 ∼2.0) and the Agilent Bioanalyzer (RIN > 8). RNA-seq libraries were prepared using poly(A) enrichment (Novogene, Cambridge; Glasgow MVLS Shared Research Facilities). Reads were aligned to the GRCm38 mouse genome using HISAT2 v2.1.0, and gene counts were obtained with featureCounts v1.4.6. Gene IDs were converted in R using BiocManager. Differential expression was analysed using DESeq2 with Padj < 0.01 and |log₂FC| > 2 as significance thresholds. Principal component analysis was performed using DEBrowser ^29^.

### Mass spectrometry glycomics

Glycan extraction from gut tissue was performed using a protocol from ^30^. Briefly, tissue was open longitudinally, washed in cold PBS and extensively homogenised on ice in lysis buffer (25 mM Tris, 150 mM NaCl, 5 mM EDTA, 1% CHAPS). Glycosphingolipids were removed by methanol/chloroform extraction. Protein pellets were reduced with DTT, carboxymethylated with iodoacetic acid, digested with trypsin, dialysed overnight against 50 mM ammonium bicarbonate, and lyophilised to generate glycopeptides. Glycopeptides were further digested with trypsin, purified on C18 cartridges, and treated with PNGase F to remove N-glycans. O-glycans were released by reductive β-elimination of the remaining glycopeptides (0.1 M KBH₄), desalted on Dowex 50W X8, derivatised, permethylated, purified on Sep-Pak C18, and lyophilised. Permethylated O-glycans were subjected to MALDI-TOF and MS/MS spectra were acquired using an AB 4800 Plus mass spectrometer. Glycan structures were assigned and annotated using GlycoWorkbench.

### 3D in vitro cultures

Human intestinal fibroblasts (CCD-18Co), epithelial cells (Caco-2), and goblet cells (HT-29) cell lines were cultured in DMEM containing 10% FCS, 0.1 mM non-essential amino acids, 2 mM L-glutamine, and 100 U/mL penicillin/streptomycin. For generation of conditioned medium by fibroblasts, cells were seeding in 2D cultures and stimulated with TNF (10 ng/ml) for 12 hours. 3D Alvetex scaffolds (Reprocell) were overlaid with 2 mg/mL rat tail collagen I for 3 h, incubated overnight, and seeded with a 9:1 mixture of Caco-2:HT-29 cells (0.5 × 10□ cells/well). After attachment (3–4 h), cultures were topped with fresh medium and maintained for downstream assays. Barrier permeability was assessed by adding HRP (100 μg/mL) to the upper chamber. After 1 h, medium was collected from the lower chamber, HRP activity was detected using TMB substrate, stopped with 10% H₂SO₄, and absorbance was measured at 450 nm (Zhu et al., 2012).

### Statistical analysis

Statistical analyses were performed using GraphPad Prism 8. Data distribution was assessed before testing. One-way ANOVA was used for comparisons involving more than two groups, and Student’s t-test or Mann–Whitney U test was used for pairwise comparisons depending on normality. Significance thresholds were p<0.05, p<0.01, p<0.001, and p<0.0001; non-significance is indicated as ns. Exact tests used for each dataset and the number of independent replicates is specified in the figure legends.

## Discussion

Despite extensive research, the mechanisms driving persistent inflammation and chronic disease progression in inflammatory arthritis remain incompletely understood. While joint⍰intrinsic processes clearly contribute to disease, increasing evidence implicates extra⍰articular tissues, including the gut, in shaping systemic inflammatory responses. Changes in intestinal permeability and stromal or epithelial activation have been proposed as components of the gut-joint axis. However, how systemic inflammation is reflected at the level of epithelial biosynthetic programs in the gut remains poorly defined. Mucin O⍰glycosylation represents a highly regulated epithelial output, integrating cellular differentiation state and inflammatory signals into the composition of secreted and cell⍰associated mucins. The molecular pathways identified in this work, describing how epithelial O⍰glycosylation is dysregulated in arthritis, provides critical insight into how systemic inflammation alters gut epithelial biology and establishes a molecular framework needed for future functional and translational research. Future research may address the functional consequences of altered O⍰glycans on epithelial barrier properties, such as permeability, something that is not directly assessed in this study.

We describe here that systemic inflammation remodels epithelial O-glycosylation, through a mechanism that may involve fibroblast paracrine contribution. Using a combination of lectin-based histochemistry, transcriptional analysis and mass spectrometry based glycomics, we show that mice undergoing experimental arthritis exhibit a marked reduction in fucosylated O-glycans in the colonic epithelium, but not in the ileum. This reduction in terminal fucosylation was consistently observed by multiple independent approaches and occurred without a corresponding decrease in epithelial cell numbers, indicating that it reflects altered epithelial glycome rather than loss of epithelial mass. To compare health versus arthritis, we chose modern omics approaches, transcriptomics and MS-based glycomics, rather than more traditional lectin-based techniques. Lectin staining of gut sections provides valuable in situ insight into the distribution and relative enrichment of specific glycan epitopes within the MUC2-positive mucus, but it is inherently limited in specificity. For example, lectins are commonly able to bind to secondary motifs with lower affinity. Furthermore, regarding specific glycosylation of the gut epithelium, lectin binding does not exclusively target epithelial cells and MUC2 associated glycans, as they also bind other cells present within or adjacent to the mucus layer. Therefore, we looked for methods providing highly specific information to specifically target the epithelial compartment. Overall, the generated results from the murine model provide a strong correlation between O-glycan remodelling and arthritic conditions, but whether loss of fucosylation is a disease driver, perpetuator, or a result of ongoing inflammation is still unclear. To further advance on the understanding of the molecular mechanisms, we used 3D in vitro models to prove that loss of epithelial fucosylation occurs in response to inflammatory pathways. These in vivo and in vitro data suggest that systemic inflammatory stress in arthritis is accompanied by selective reprogramming of epithelial glycosylation pathways.

Our in vitro work provides initial data towards understanding the cellular mechanisms responsible for changes in epithelial fucosylation, directing future research in this area towards stromal-epithelial interactions. Although TNF did not appear to have a direct effect on epithelial glycosylation, mediation of fibroblasts in this process mimicked our observations in vivo. Fibroblasts were initially identified as matrix regulators. In the gut, they produce collagen type I, II, and V, and fibronectin, and matrix remodelling MMPs ^31^. However, their functions in immunity and epithelial homeostasis are being increasingly recognised ^32, 33^. Intestinal fibroblasts control tissue repair mediated by Gremlin-1 and R-spondin-1, gut immunity [secretion of IL-7, Ccl2, Ptgs2, amphiregulin, and maintenance of the epithelial crypt environment [Wnt2/Wnt2b, BMP antagonists] ^34, 35^. These functions are associated with functional and anatomical specialization, since fibroblasts in the distal villi areas secrete BMP3/7 to counteract WNT signalling, contributing to epithelial cell differentiation. Our *in vitro* results confirm that a fibroblast-derived secretome upon TNF activation affects epithelial O-glycan fucosylation. It is possible that *in vivo*, only PDGFRα^hi^ fibroblasts mediate O-glycan regulation, due to their function in BMP-induced epithelial differentiation, but further *in vivo* work is required to test this hypothesis. Nevertheless, our 3D model provides initial experimental support, contributing to design of more targeted experiments to determine whether mucin O-glycosylation is directly involved in gut barrier integrity. This model mimics TNF-mediated inflammatory responses relevant for the gut environment, such as regulation of epithelial barrier permeability and mucus synthesis. In recent years, there has been an increase in research efforts dedicated to the development of functional three-dimensional gut models and organ-on-a-chip devices. There is no doubt that these models will contribute to understand fibroblast-epithelial crosstalk, and the incorporation of modelling physiological mucin glycosylation should be a priority to advance the field.

Changes in epithelial O-glycosylation, perhaps driven by a TNF-fibroblast-fucosylation axis, could have major consequences in gut physiology, and perpetuation of inflammation in arthritis. Our results suggests that the main driver in glycan remodelling is the reduction of terminally fucosylated O-glycans, supported by mass spectrometry data and transcriptomic profile, indicating a strong down-regulation of FUT2 and FUT4. Differences in mucin fucosylation could weaken epithelial barrier integrity, and can also alter the microbial composition ^36 37^, which in turn can amplify systemic inflammation. All these alterations could potentially contribute to joint pathology, thereby exacerbating chronic disease. In our arthritis model, colonic epithelial cells show a strong down-regulation of FUT2, enzyme responsible of mucin terminal fucosylation. Tang et al showed that down-regulation of FUT2 is associated in IBD patients and mouse models of colitis, with knocked-out FUT2 mice showing increased susceptibility to inflammation ^37^. Overall, there is an increasing support for the pathogenic role of fucosylation in intestinal inflammation, both locally, and also systemically as reported here.

We have previously described a loss of microbial diversity in the colon of Collagen-Induced Arthritis (CIA) mice ^21^, in accordance with the widely accepted correlation between the gut microbiome and both local and systemic inflammation, including rheumatoid arthritis. O-glycans are essential to maintain healthy microbiomes in the gut, since many commensal bacterial species forage on them and contribute to their turnover. For example, Bacteroides in general can metabolize host fucosylated-glycans, supporting microbial stability under nutrient-limited conditions ^38^, and fucosidases are also expressed by Bifidobacterium ^39^. Interestingly, host mucin fucosylation is also reinforced by microbiome-derived signals, creating protective feedback loops. Aryl hydrocarbon receptor (AhR) ligands and butyrate are bacterial metabolites known to protect against arthritis ^40^ and up-regulate FUT2 expression ^41^. These positive feedback loops would be eliminated during arthritis, contributing to disease perpetuation. Since glycans are central in host-microbiome feedback, therapies targeting, or restoring, specific glycan epitopes could be an alternative for clinical interventions in inflammatory arthritis. Besides microbiome-related work, our study suggests other directions to follow, although it contains limitations. As mentioned before, *in vivo* causality experiments are needed prior to any clinical or preclinical study. We show evidence of the existence of a fibroblast-epithelial fucosylation axis that is triggered by TNF activation in vitro. Likewise, we observed a relative increase in sialic acid containing O-glycans, but in this case we do not have the same level of certainty to establish mechanistic pathways, since transcriptomics and glycomics data do not fully correlate. This is not unusual in the glycobiology field, since transcriptomics alone are insufficient to predict actual glycan expression given the non-lineal nature of glycan synthesis. The relative increase of sialylated structures could be a consequence of the reduced fucosylation. Furthermore, analysis of O-glycans relies on chemical reactions, such as reductive elimination. This approach is less clean than N-glycan analysis, which uses a specific enzyme to cleave sugars from glycoproteins. Therefore, although O-glycan analysis gave us enough information to identify the main core structures by MS, motifs with higher molecular weight could not be included in the analysis due to the increased signal/threshold ratio. We could not confirm the presence of increased sialylated forms, which fall in larger structures with bigger m/z values. For translational purposes, restoration of fucosylation by dietary supplementation or genetic approaches would provide further mechanistic support, as well as fibroblast-specific inhibition. Likewise, integration with microbiome sequencing would confirm the involvement of mucin fucosylation in regulating pathogenic networks in arthritis. A combined approach of in vivo and in vitro methods would be required to provide conclusive causality in the field. This is an important challenge in the context of the gut-joint axis theory, due to the difficulty of conducting pre-clinical work in arthritic patients.

Mucin glycosylation is a very relevant area of research to identify new therapeutic targets in the gut, and more work is expected to fill current gaps in our understanding of this interesting organ. Our work identifies a previously unrecognised axis: TNF-driven fibroblast activation leading to epithelial glycosylation remodelling. Together, these results indicate that fibroblast activation can influence epithelial glycan presentation, providing a potential mechanistic link between stromal inflammation and epithelial glycosylation changes observed during arthritis.

## Acknowledgments

This work was funded by an Industrial PhD studentship to P.P. from the University of Glasgow and a Career Development award to M.A.P. from Versus Arthritis (21221).

---

**Supplementary Figure 1.**
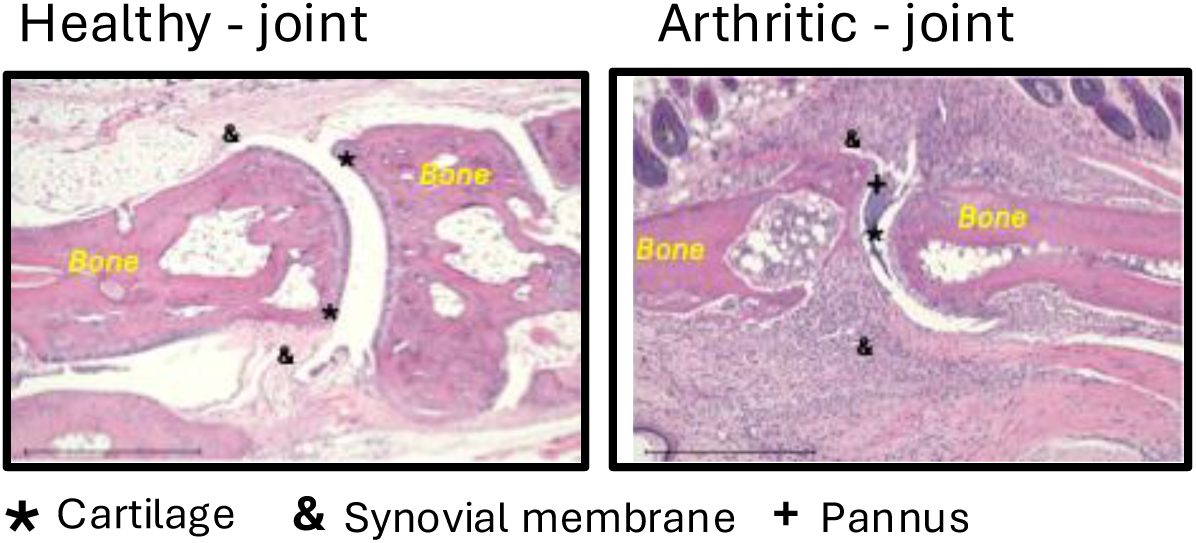
Histological assessment of joint architecture in naïve and arthritic mice. Representative histological sections of joints from naïve (left) and arthritic (right) mice. In naïve joints, normal tissue architecture is preserved, with intact bone surfaces and well-defined joint spaces. In contrast, arthritic joints show marked pathological changes, including synovial hyperplasia, inflammatory cell infiltration, and disruption of joint integrity. Bone regions are indicated (Bone), while symbols denote key pathological features: (*) highlights areas of tissue damage or erosion, (&) indicates synovial expansion or inflammatory infiltrate, and (+) marks regions of pronounced joint space alteration. Scale bars as indicated.

**Supplementary figure 2.**
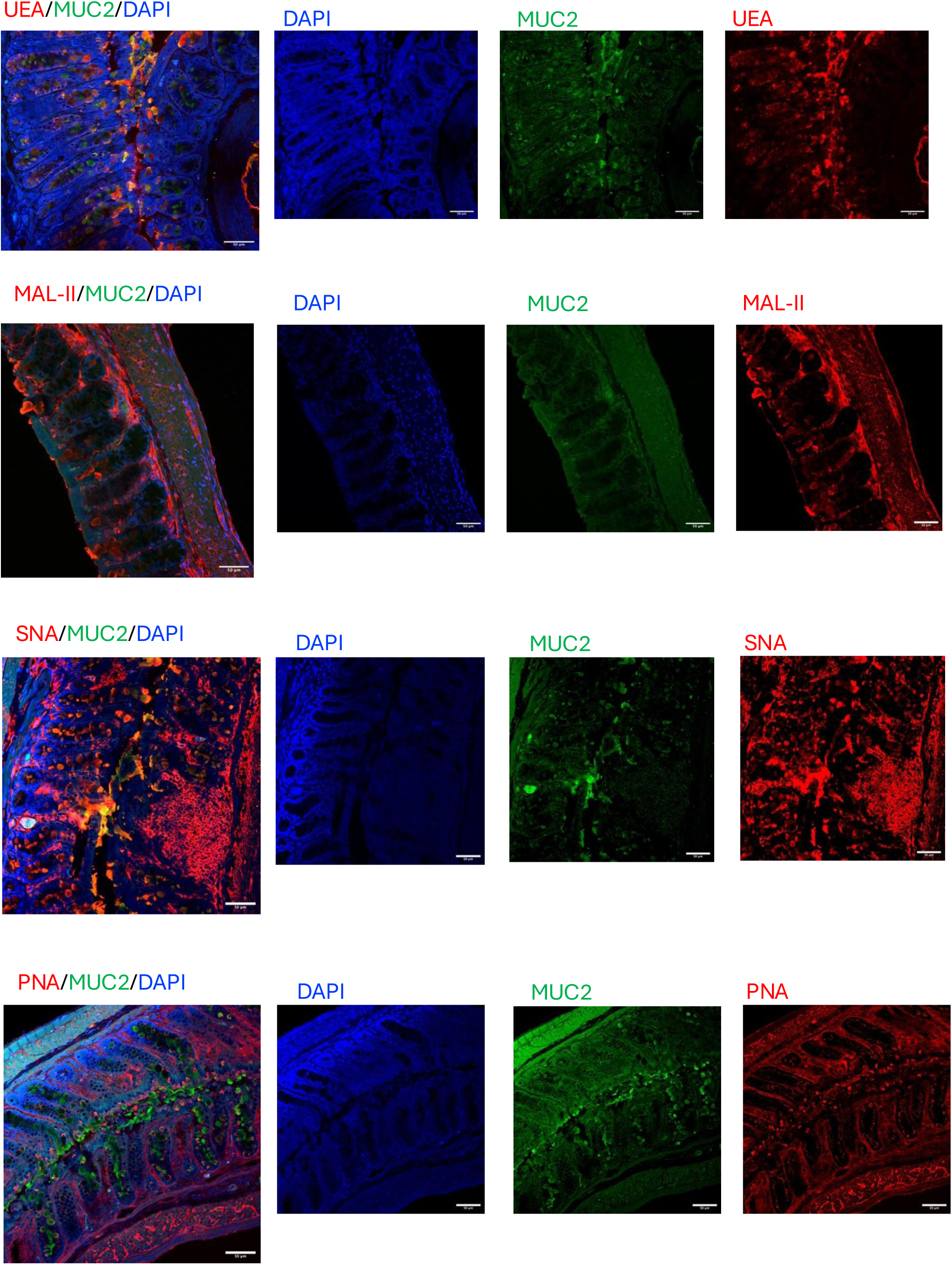
Lectin staining reveals glycan heterogeneity within the colonic mucus barrier of naïve mice. Representative confocal immunofluorescence images of colonic sections from naïve mice co⍰stained with lectins and the mucus marker MUC2. Sections were stained with Ulex europaeus agglutinin I (UEA⍰I; α1,2⍰fucosylated glycans), Maackia amurensis lectin II (MAL⍰II; α2,3⍰sialylated glycans), Sambucus nigra agglutinin (SNA; α2,6⍰sialylated glycans), or peanut agglutinin (PNA; Galβ1⍰3GalNAc) [Lectins in red] together with anti MUC2 antibody (green) to delineate the mucus layer and mucus-producing cells. Nuclei are counterstained with DAPI (Blue). Merged images and individual channels are shown, demonstrating distinct spatial patterns of lectin binding within the MUC2⍰positive mucus layer and colonic epithelium. Scale bars, 50 µm.

## References

1. Pelaseyed T, Hansson GC. Membrane mucins of the intestine at a glance. J Cell Sci 2020; 133(5).

2. Johansson ME, Larsson JM, Hansson GC. The two mucus layers of colon are organized by the MUC2 mucin, whereas the outer layer is a legislator of host-microbial interactions. Proc Natl Acad Sci U S A 2011; 108 **Suppl 1**(Suppl 1): 4659–4665.

3. Bennett EP, Mandel U, Clausen H, Gerken TA, Fritz TA, Tabak LA. Control of mucin-type O-glycosylation: a classification of the polypeptide GalNAc-transferase gene family. Glycobiology 2012; 22(6): 736–756.

4. Gill DJ, Clausen H, Bard F. Location, location, location: new insights into O-GalNAc protein glycosylation. Trends Cell Biol 2011; 21(3): 149–158.

5. Robbe C, Capon C, Maes E, Rousset M, Zweibaum A, Zanetta JP et al. Evidence of regio-specific glycosylation in human intestinal mucins: presence of an acidic gradient along the intestinal tract. J Biol Chem 2003; 278(47): 46337–46348.

6. Robbe C, Capon C, Coddeville B, Michalski JC. Structural diversity and specific distribution of O-glycans in normal human mucins along the intestinal tract. Biochem J 2004; 384(Pt 2): 307–316.

7. Holmen Larsson JM, Thomsson KA, Rodriguez-Pineiro AM, Karlsson H, Hansson GC. Studies of mucus in mouse stomach, small intestine, and colon. III. Gastrointestinal Muc5ac and Muc2 mucin O-glycan patterns reveal a regiospecific distribution. Am J Physiol Gastrointest Liver Physiol 2013; 305(5): G357–363.

8. Pickard JM, Maurice CF, Kinnebrew MA, Abt MC, Schenten D, Golovkina TV et al. Rapid fucosylation of intestinal epithelium sustains host-commensal symbiosis in sickness. Nature 2014; 514(7524): 638–641.

9. Goto Y, Obata T, Kunisawa J, Sato S, Ivanov, II, Lamichhane A et al. Innate lymphoid cells regulate intestinal epithelial cell glycosylation. Science 2014; 345(6202): 1254009.

10. Turroni F, Milani C, Duranti S, Mahony J, van Sinderen D, Ventura M. Glycan Utilization and Cross-Feeding Activities by Bifidobacteria. Trends Microbiol 2018; 26(4): 339–350.

11. Chang J, Leong RW, Wasinger VC, Ip M, Yang M, Phan TG. Impaired Intestinal Permeability Contributes to Ongoing Bowel Symptoms in Patients With Inflammatory Bowel Disease and Mucosal Healing. Gastroenterology 2017; 153(3): 723–731 e721.

12. van der Post S, Jabbar KS, Birchenough G, Arike L, Akhtar N, Sjovall H et al. Structural weakening of the colonic mucus barrier is an early event in ulcerative colitis pathogenesis. Gut 2019; 68(12): 2142–2151.

13. Larsson JM, Karlsson H, Crespo JG, Johansson ME, Eklund L, Sjovall H et al. Altered O-glycosylation profile of MUC2 mucin occurs in active ulcerative colitis and is associated with increased inflammation. Inflamm Bowel Dis 2011; 17(11): 2299–2307.

14. Wei J, Chen C, Feng J, Zhou S, Feng X, Yang Z et al. Muc2 mucin O-glycosylation interacts with enteropathogenic Escherichia coli to influence the development of ulcerative colitis based on the NF-kB signaling pathway. J Transl Med 2023; 21(1): 793.

15. McGovern DP, Jones MR, Taylor KD, Marciante K, Yan X, Dubinsky M et al. Fucosyltransferase 2 (FUT2) non-secretor status is associated with Crohn’s disease. Hum Mol Genet 2010; 19(17): 3468–3476.

16. Fu J, Wei B, Wen T, Johansson ME, Liu X, Bradford E et al. Loss of intestinal core 1-derived O-glycans causes spontaneous colitis in mice. J Clin Invest 2011; 121(4): 1657–1666.

17. Wang Y, Huang D, Chen KY, Cui M, Wang W, Huang X et al. Fucosylation Deficiency in Mice Leads to Colitis and Adenocarcinoma. Gastroenterology 2017; 152(1): 193–205 e110.

18. Pacheco AR, Curtis MM, Ritchie JM, Munera D, Waldor MK, Moreira CG et al. Fucose sensing regulates bacterial intestinal colonization. Nature 2012; 492(7427): 113–117.

19. Ke J, Li Y, Han C, He R, Lin R, Qian W et al. Fucose Ameliorate Intestinal Inflammation Through Modulating the Crosstalk Between Bile Acids and Gut Microbiota in a Chronic Colitis Murine Model. Inflamm Bowel Dis 2020; 26(6): 863–873.

20. Matei DE, Menon M, Alber DG, Smith AM, Nedjat-Shokouhi B, Fasano A, et al. Intestinal barrier dysfunction plays an integral role in arthritis pathology and can be targeted to ameliorate disease. Med (N Y) 2021; 2(7): 864–883 e869.

21. Pan P, Wang Y, Nyirenda MH, Saiyed Z, Karimian Azari E, Sunderman A et al. Undenatured type II collagen protects against collagen-induced arthritis by restoring gut-joint homeostasis and immunity. Commun Biol 2024; 7(1): 804.

22. Doonan J, Tarafdar A, Pineda MA, Lumb FE, Crowe J, Khan AM et al. The parasitic worm product ES-62 normalises the gut microbiota bone marrow axis in inflammatory arthritis. Nat Commun 2019; 10(1): 1554.

23. Rooney CM, Jeffery IB, Mankia K, Wilcox MH, Emery P. Dynamics of the gut microbiome in individuals at risk of rheumatoid arthritis: a cross-sectional and longitudinal observational study. Ann Rheum Dis 2024.

24. Thompson KN, Bonham KS, Ilott NE, Britton GJ, Colmenero P, Bullers SJ et al. Alterations in the gut microbiome implicate key taxa and metabolic pathways across inflammatory arthritis phenotypes. Sci Transl Med 2023; 15(706): eabn4722.

25. Zaiss MM, Joyce Wu HJ, Mauro D, Schett G, Ciccia F. The gut-joint axis in rheumatoid arthritis. Nat Rev Rheumatol 2021; 17(4): 224–237.

26. Pineda MA, McGrath MA, Smith PC, Al-Riyami L, Rzepecka J, Gracie JA et al. The parasitic helminth product ES-62 suppresses pathogenesis in collagen-induced arthritis by targeting the interleukin-17-producing cellular network at multiple sites. Arthritis Rheum 2012; 64(10): 3168–3178.

27. Davidson S, Coles M, Thomas T, Kollias G, Ludewig B, Turley S et al. Fibroblasts as immune regulators in infection, inflammation and cancer. Nat Rev Immunol 2021; 21(11): 704–717.

28. Erben U, Loddenkemper C, Doerfel K, Spieckermann S, Haller D, Heimesaat MM et al. A guide to histomorphological evaluation of intestinal inflammation in mouse models. Int J Clin Exp Pathol 2014; 7(8): 4557–4576.

29. Kucukural A, Yukselen O, Ozata DM, Moore MJ, Garber M. DEBrowser: interactive differential expression analysis and visualization tool for count data. 2019: 1–12.

30. Ismail MN, Stone EL, Panico M, Lee SH, Luu Y, Ramirez K et al. High-sensitivity O-glycomic analysis of mice deficient in core 2 beta1,6-N-acetylglucosaminyltransferases. Glycobiology 2011; 21(1): 82–98.

31. Bonnans C, Chou J, Werb Z. Remodelling the extracellular matrix in development and disease. Nat Rev Mol Cell Biol 2014; 15(12): 786–801.

32. Brugger MD, Basler K. The diverse nature of intestinal fibroblasts in development, homeostasis, and disease. Trends Cell Biol 2023; 33(10): 834–849.

33. Chalkidi N, Paraskeva C, Koliaraki V. Fibroblasts in intestinal homeostasis, damage, and repair. Front Immunol 2022; 13: 924866.

34. Stzepourginski I, Nigro G, Jacob JM, Dulauroy S, Sansonetti PJ, Eberl G et al. CD34+ mesenchymal cells are a major component of the intestinal stem cells niche at homeostasis and after injury. Proc Natl Acad Sci U S A 2017; 114(4): E506–E513.

35. McCarthy N, Manieri E, Storm EE, Saadatpour A, Luoma AM, Kapoor VN et al. Distinct Mesenchymal Cell Populations Generate the Essential Intestinal BMP Signaling Gradient. Cell Stem Cell 2020; 26(3): 391–402 e395.

36. Cheng S, Hu J, Wu X, Pan JA, Jiao N, Li Y et al. Altered gut microbiome in FUT2 loss-of-function mutants in support of personalized medicine for inflammatory bowel diseases. J Genet Genomics 2021; 48(9): 771–780.

37. Tang X, Wang W, Hong G, Duan C, Zhu S, Tian Y et al. Gut microbiota-mediated lysophosphatidylcholine generation promotes colitis in intestinal epithelium-specific Fut2 deficiency. J Biomed Sci 2021; 28(1): 20.

38. van der Post S, Subramani DB, Backstrom M, Johansson MEV, Vester-Christensen MB, Mandel U et al. Site-specific O-glycosylation on the MUC2 mucin protein inhibits cleavage by the Porphyromonas gingivalis secreted cysteine protease (RgpB). J Biol Chem 2013; 288(20): 14636–14646.

39. Ashida H, Miyake A, Kiyohara M, Wada J, Yoshida E, Kumagai H et al. Two distinct alpha-L-fucosidases from Bifidobacterium bifidum are essential for the utilization of fucosylated milk oligosaccharides and glycoconjugates. Glycobiology 2009; 19(9): 1010–1017.

40. Rosser EC, Piper CJM, Matei DE, Blair PA, Rendeiro AF, Orford M et al. Microbiota-Derived Metabolites Suppress Arthritis by Amplifying Aryl-Hydrocarbon Receptor Activation in Regulatory B Cells. Cell Metab 2020; 31(4): 837–851 e810.

41. Zhang DD, Huang ZX, Liu XC, Ding XP, Li L, He Y et al. Butyrate protects the intestinal barrier by upregulating Fut2 expression via MEK4-JNK signaling pathway activation. Pediatr Res 2025; 97(1): 128–137.

