## Supplementary figures for "Arthritis-Associated Inflammation Remodels Colonic O-Glycosylation"

Healthy - joint    Arthritic - joint

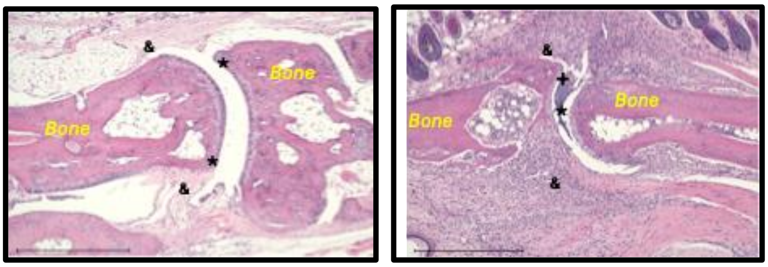


★ Cartilage   & Synovial membrane   + Pannus

Supplementary Figure 1

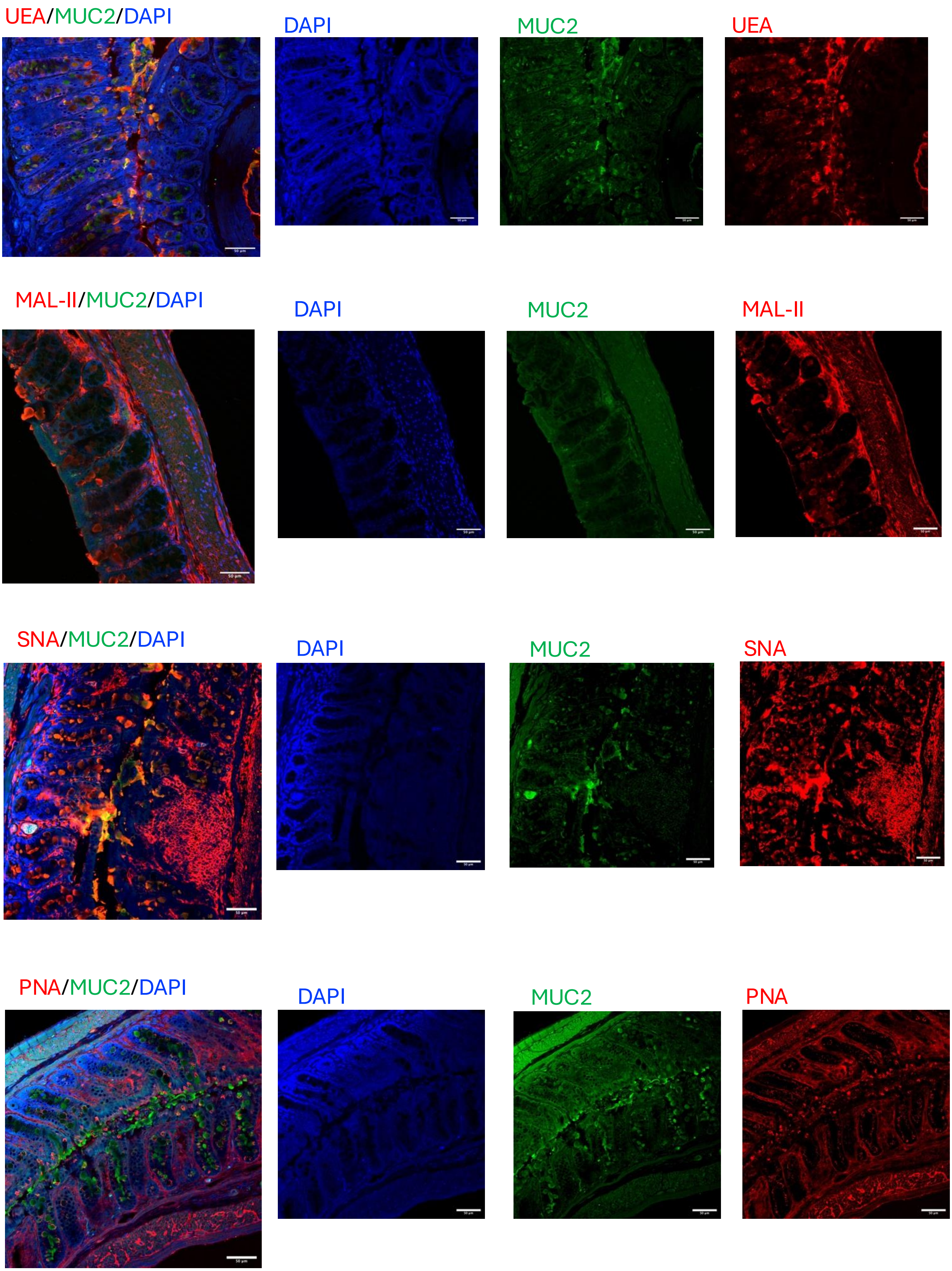


Supplementary Figure 2
